# Simple models of quantitative firing phenotypes in hippocampal neurons: comprehensive coverage of intrinsic diversity

**DOI:** 10.1101/632430

**Authors:** Siva Venkadesh, Alexander O. Komendantov, Diek W. Wheeler, David J. Hamilton, Giorgio A. Ascoli

## Abstract

Patterns of periodic voltage spikes elicited by a neuron help define its dynamical identity. Experimentally recorded spike trains from various neurons show qualitatively distinguishable features such as delayed spiking, spiking with/without frequency adaptation, and intrinsic bursting. Moreover, the input-dependent responses of a neuron not only show different quantitative features, such as higher spike frequency for a stronger input current injection, but can also exhibit qualitatively different responses, such as spiking and bursting under different input conditions, thus forming a complex phenotype of responses. In a previous work, *Hippocampome.org*, a comprehensive knowledgebase of hippocampal neuron types, systematically characterized various spike pattern phenotypes experimentally identified from 120 neuron types/subtypes. In this paper, we present a comprehensive set of simple phenomenological models that quantitatively reproduce the diverse and complex phenotypes of hippocampal neurons. In addition to point-neuron models, we created compact multi-compartment models with up to four compartments, which will allow spatial segregation of synaptic integration in network simulations. Electrotonic compartmentalization observed in our compact multi-compartment models is qualitatively consistent with experimental observations. Furthermore, we observed that adding dendritic compartments to point-neuron models, in general, allowed soma to reproduce features of bursting patterns and abrupt non-linearities in some frequency adapting patterns slightly more accurately. This work maps 120 neuron types/subtypes in the rodent hippocampus to a low-dimensional model space and adds another dimension to the knowledge accumulated in *Hippocampome.org*. Computationally efficient representations of intrinsic dynamics, along with other pieces of knowledge available in *Hippocampome.org*, provide a biologically realistic platform to explore the dynamical interactions of various types at the mesoscopic level.

**Author Summary:** The neurons in the hippocampus show enormous diversity in their intrinsic activity patterns. A comprehensive characterization of various intrinsic types using a neuronal modeling system is necessary to simulate biologically realistic networks of brain regions. Morphologically detailed neuronal modeling frameworks often limit the scalability of such network simulations due to the specification of hundreds of rules governing each neuron’s intrinsic dynamics. In this work, we have accomplished a comprehensive mapping of experimentally identified intrinsic dynamics in a simple modeling system with only two governing rules. We have created over a hundred point-neuron models that reflect the intrinsic differences among the hippocampal neuron types both qualitatively and quantitatively. In addition, we compactly extended our point-neurons to include up to four compartments, which will allow anatomically finer-grained connections among the neurons in a network. Our compact model representations, which are freely available in *Hippocampome.org*, will allow future researchers to investigate dynamical interactions among various intrinsic types and emergent integrative properties using scalable, yet biologically realistic network simulations.

## 1. Introduction

Complex interactions among a myriad of neurons make it challenging to study the functions of brain regions. Although each neuron is different, their landmark features such as the dendritic structure and patterns of somatic voltage spikes help define types of neurons, and, such grouping allows for a tractable description and investigation of complex interactions in a network. For instance, large-scale network models of brain regions can include precisely defined neuronal types to create a biologically realistic platform for hypothesis testing. While neurons differ in their morphological, biochemical and electrophysiological features, precisely what features are useful and relevant for neuronal grouping is a topic of great interest [1].

A few studies have created large-scale network models of brain regions [2–5]. The major methodological difference among these studies is the level of biological details captured in the individual components of the network and there is often a tradeoff between such biological details and the scale of the network. For example, a microcircuit model of the rat somatosensory cortex [4] simulated ~31,000 neurons with ~37 million synapses, where each neuron was a biophysically detailed description of one of 207 morpho-electrical types identified experimentally. On the other hand, a large-scale description of thalamocortical systems [2], which used simplified phenomenological neuron models [6], simulated a network of much larger scale (one million neurons and half a billion synapses), but it only included 22 abstract types among the neurons. In current work, with a vision of creating a real-scale network model of the rodent hippocampus that nevertheless captures biological details at the mesoscopic level, we have created phenomenological models of 120 hippocampal neuron types and subtypes using their intrinsic dynamics identified experimentally. More recently, a large-scale effort [7] created a database of simple models for hundreds of neurons of various transgenic types in the mouse primary visual cortex with a similar vision.

A large-scale literature mining effort created Hippocampome.org [8], a comprehensive knowledgebase of neuron types in rodent hippocampal formation (dentate gyrus, CA3, CA2, CA1, subiculum, and entorhinal cortex). It provides information on morphology, electrophysiology, and molecular marker profiles of more than 100 neuron types, where the type of a neuron is primarily determined based on its neurite invasion pattern across hippocampal parcels. Latest enhancement to this knowledge base annotated 90 of these morphological types with their spike patterns and identified a total of 120 neuron types/subtypes [9]. Features of experimentally recorded spike patterns were extracted for a neuron type from relevant publications and a systematic characterization of spike pattern features revealed nine unique families of intrinsic dynamics such as delayed spiking, non-adapting spiking, simple adapting spiking, and persistent stuttering among hippocampal neurons. Furthermore, many neuron types exhibit different classes of spike patterns for different input currents resulting in complex spike pattern phenotypes.

In this article, we present a comprehensive set of point neuron models that quantitatively reproduce various spike pattern phenotypes of hippocampal neurons. We also created multi-compartment models that are compact extensions of point neurons in order to allow spatial context for synaptic integration in a network. In addition, our compact multi-compartment models exhibit electrotonic properties consistent with experimental observations. We also report interesting relationships between the abstract model parameters and various biological properties. The models were created using an automated modeling framework [10], and they further enhance the existing accumulated knowledge in *Hippocampome.org*, where they are freely available to download. By identifying several possibilities for a quantitative phenotype in phenomenological space, current work comprehensively maps hippocampal neuron types to low-dimensional model subspaces, which can be used as sampling regions for biologically realistic large-scale network simulations of hippocampal circuits.

## 2. Methods

The class of a spike pattern is identified based on various transient and/or steady-state elements present in the pattern. Transient elements are Delay (D), if the first spike latency (*fsl*) is sufficiently long; Adapting Spiking (ASP), if the inter-spike intervals (*ISIs*) increase over time showing a spike frequency adaptation (*sfa*); Rapidly Adapting Spiking (RASP), if a strong *sfa* is only present in the first two or three ISIs, Transient Stuttering (TSTUT), if a quiescent period follows a cluster of high frequency spikes; and Transient Slow-Wave Bursting (TSWB), if a slow after-hyperpolarizing potential follows a cluster of high frequency spikes. Steady-state elements are Silence (SLN), if the post-spike silence (pss) (quiescence following the last spike) is sufficiently long; Non-Adapting Spiking (NASP), if no frequency adaptation is identified in a non-interrupted spiking; Persistent Stuttering (PSTUT), if at least one sufficiently long quiescent period separates two clusters of high frequency spikes; and Persistent Slow-Wave Bursting (PSWB) if a slow after-hyperpolarizing potential is present in an otherwise PSTUT pattern. Thus, the key features are *fsl, sfa, pss* and the number of ISIs (*nISIs*) for a spiking pattern, and burst widths (*bw*), post-burst intervals (*pbi*), number of bursts (*n_bursts*) and *nISIs* within a burst (*b-nISIs*) for a stuttering/bursting pattern. Refer to [9] for more details on the criteria for various spike pattern classes. These temporal features identify the class of a single spike pattern, and all classes of patterns exhibited by a neuron under different input currents collectively define the spike pattern phenotype of that neuron. Thus, our approach emphasizes the temporal patterns in the periodic voltage spikes rather than the shape of the spike or subthreshold dynamics to define the intrinsic dynamics.

We used a two-dimensional quadratic model (QM) [6, 11] to reproduce the spike pattern phenotypes. This model is governed by the state variables membrane voltage (V) and membrane recovery variable (U):

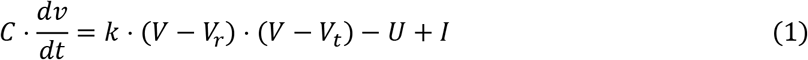

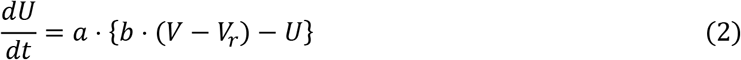

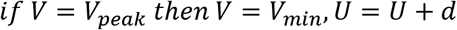

where *V_r_* and *V_t_* are the resting and threshold voltages respectively. *V_peak_* is the spike cutoff value, *V_min_* is the post-spike reset value for the voltage and *C* is the cell capacitance. The parameters *k, a, b* and *d* affect the model’s intrinsic dynamics both qualitatively (e.g. the type of bifurcation revealed by fast-spiking and non-fast-spiking behaviors) and quantitatively (e.g. excitability level and magnitude of *sfa*). Compact multi-compartment models with up to four compartments were modeled using an asymmetric coupling mechanism by calculating coupling currents in somatic (*I_soma_*) and dendritic (*I_dend_*)compartments as follows:

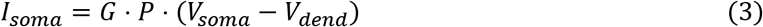

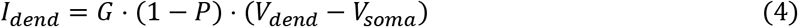

Where *G* is the coupling strength and *P* denotes the degree of coupling asymmetry, which determines the influence of a compartment on the overall model dynamics [12]. As reported before [10], most of our compact multi-compartment models specify a much weaker coupling toward the soma than away from it, making the somatic compartment dominate the overall model intrinsic dynamics. Compact multi-compartment models were also constrained to exhibit appropriate relative excitabilities and input resistances between soma and dendrites, and sub- and supra-threshold signal propagation properties.

Our modeling framework [10] uses evolutionary algorithms (EA) and employs a feature-based error function. By incorporating spike pattern features (*fsl, sfa* etc.) and qualitative class criteria (*delay factor, number of piecewise linear fit parameters of ISIs* etc.) [9] in the error landscape, our approach enforces a fine level of granularity in the key quantitative features of various spike pattern classes. The operators of the EA (mutation, crossover etc.) were configured by taking into account the features of error landscape created by the QM parameters [13]. In order for a model found by the EA to be accepted, the classes of its spike patterns must match those of experimental traces. There is, however, one exception: Without additional dendritic dynamics, the QM failed to reproduce RASP.ASP. class of patterns, which show a strong and rapid adaptation (in the first 2 or 3 ISIs) followed by a very weak and sustained adaptation. Therefore, single-compartment models of seven neuron types, which experimentally showed this complex transient pattern, were accepted with RASP.NASP patterns instead (see results).

Pairwise correlations were performed to explore the relationships between QM parameters and various pieces of knowledge such as biomarker expression that have been accumulated in Hippocampome.org. Continuous QM parameters were converted into categorical variables appropriately by marking positive and negative or by labelling top- and bottom-one-third ranges respectively as high and low. Correlations between the categorical variables were evaluated using Barnard’s exact test for 2×2 contingency tables. This test provides the greatest statistical power when row and column totals are free to vary [14].

## 3. Results

### 3.1. Single-compartment models of diverse intrinsic spike pattern phenotypes

The intrinsic dynamics of a neuron is identified in experiments typically by injecting step input currents of various magnitudes. A neuron’s responses to these inputs typically fall into one of two phenotype super-families: (1) spiking phenotype, where the neuron only exhibits continuous spike pattern classes such as ASP.SLN, NASP, and D.NASP for different input currents (Fig 1A-D), and (2) stuttering/bursting phenotype, where the neuron exhibits an interrupted spike pattern class such as TSWB.SLN, TSTUT.NASP, and PSTUT for at least one input current (Fig 2A-D). A spiking or stuttering phenotype could be formed by various combinations of spike pattern classes, and models for four exemplar cases in each of these two phenotype super-families are reported in this article (visit *Hippocampome.org* for a comprehensive list of phenotypes and their models).

**Fig 1.**
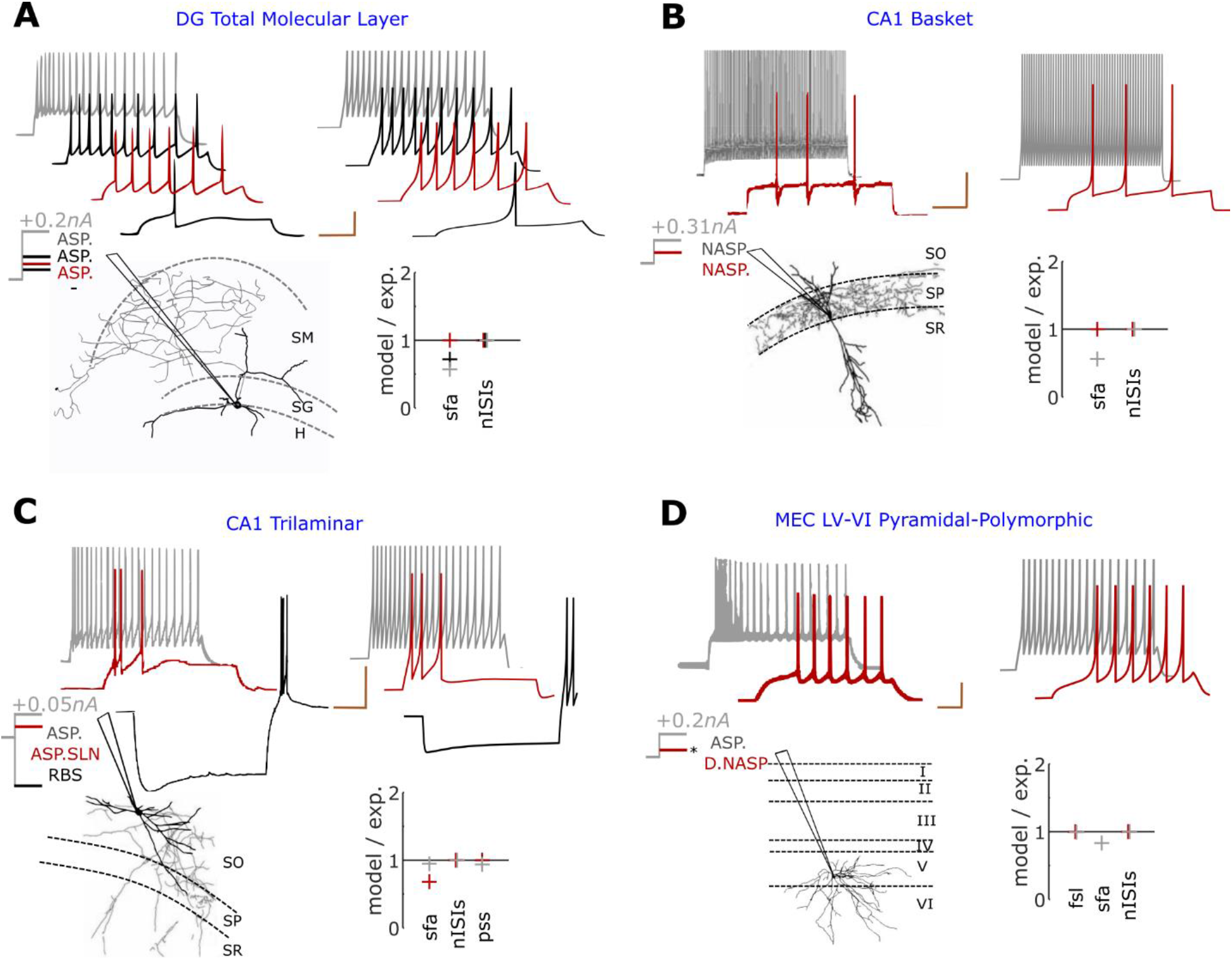
Exemplar models of continuous spiking phenotypes. In each panel, experimentally recorded voltage traces are given in top-left, and their morphological identity and magnitudes of somatic current injections are given in bottom-left. The traces were digitized by *Hippocampome.org*. Morphological abbreviations: SO – stratum oriens, SP – stratum pyramidale, SR – stratum radiatum, SO – stratum oriens, SG – stratum granulosum, H – hilus. Model responses for similar input currents (±0.01*nA* from experimental input) are given in top-right and the goodness of fit is given for key features as the ratio between simulated and experimentally recorded values in bottom-right. The traces are highlighted in different colors to visually compare experimental and model responses and to identify the input current and key features for each trace. Calibration bars denote 200ms and *20mV* in all panels. (**A**) Simple phenotype of a dentate gyrus (DG) Total Molecular Layer neuron that elicits patterns of class ASP. under three different input currents [15]. Digitally reconstructed morphology was reproduced from Neuromorpho.org [16]. (**B**) Simple phenotype of a CA1 Basket neuron that elicits patterns of class NASP for +0.15*nA* and +0.31*nA* [17]. Note that *sfa* in red trace is not statistically significant to qualify this pattern as ASP. (**C**) The phenotype of a CA1 Trilaminar neuron shows different classes of patterns for +0.025*nA* and +0.05*nA* [17]. In addition, this neuron elicits rebound spikes (RBS) for a hyperpolarizing input of −0.1*nA*. (**D**) The phenotype of a medial-entorhinal cortex (MEC) neuron shows different classes of patterns for +0.2*nA* and an unknown input (denoted by ‘*’) near its excitability level [18].

**Fig 2.**
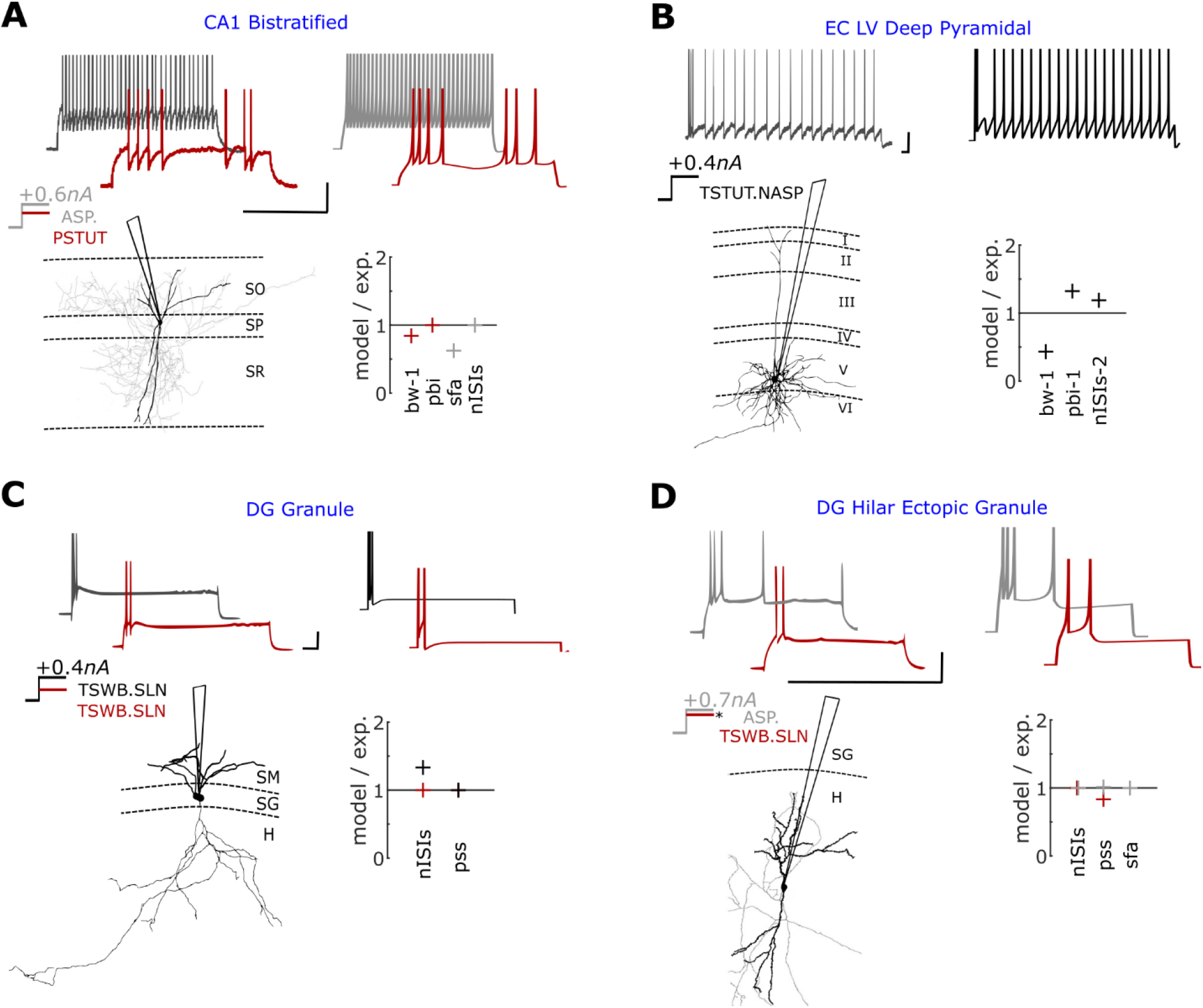
Exemplar models of stuttering/bursting phenotypes. (**A**) Complex phenotype of a Bistratified neuron in CA1. This neuron elicits a stuttering pattern for +0.4*nA* (red) and a spiking pattern for +0.6*nA* (grey) [19]. Digitally reconstructed morphology [20] was reproduced from Neuromorpho.org [16]. (**B**) The voltage trace recorded from an entorhinal layer-5 neuron shows both bursting and spiking features for +0.4*nA* [21]. (**C**) A DG granule neuron transiently bursts for both +0.2*nA* and +0.4*nA* with quantitative difference [22]. Digitally reconstructed morphology was reproduced from Neuromorpho.org [16]. (**D**) A dentate gyrus neuron that transiently bursts near its excitability level (red) elicits a spiking pattern with a strong *sfa* (grey) for a higher input current [23]. ‘*’ indicates the unknown magnitude of the input current near excitability. All voltage traces were digitized by *Hippocampome.org*. Experimental spike amplitudes are truncated. Calibration bars denote 200*ms* and 20*mV* in all panels.

In the simplest case, a neuron exhibits spike patterns of the same class regardless of the input current strength. For example, the three spike patterns recorded under different input currents from a DG Total Molecular Layer neuron were identified as ASP. (Fig 1A), and the two patterns recorded from a CA1 Basket neuron were identified as NASP. (Fig 1B). Such simple-behavior neurons typically show different quantitative features among different patterns of the same class. In the former example, the three ASP. traces were experimentally recorded under +0.075nA (red), +0.100nA (black), and +0.200nA (grey) [15]. The ISI counts (*nISIs*) are 5, 9, and 19, and *sfa* magnitudes are 0.142, 0.114, and 0.056 respectively for the red, black and grey traces. The model of this neuron type was constrained to quantitatively reproduce the spike pattern features for similar input currents: *nISIs* of 5, 9 and 19, and sfa magnitudes of 0.142, 0.082, and 0.032 respectively for +0.073*nA*, +0.102*nA* and +0.205*nA*. Note that a minimum of two spikes are required in order to identify a class, hence, single-spike traces are not assigned a class label. However, such single-spike traces help capture the excitability levels in the models more precisely.

Additionally, a neuron can show more complex behaviors by eliciting patterns of different classes under different input currents (Fig 1C-D). Both CA1 Trilaminar, and MEC LV-VI Pyramidal-Polymorphic neurons include ASP. in their phenotypes (grey traces), but they show different dynamics close to their respective excitability levels. Whereas the former quickly fired a few spikes before going into a silence mode (ASP.SLN), the latter showed delayed-spiking (D.NASP). The model quantitatively reproduces the characterizing features of these different classes (see *pss* for ASP.SLN and *fsl* for D.NASP). Also, note that the model reproduces the rebound-spiking behavior for a hyperpolarizing input current, a known feature of CA1 Trilaminar neurons [17].

Another level of complexity in spike pattern phenotypes is when the intrinsic dynamics show sharply distinguishable spike pattern classes under different input conditions. For example, a CA1 bistratified neuron stutters (PSTUT) for +0.4*nA*, and spikes for +0.6*nA* (ASP.) (Fig 2A). A few neuron types and subtypes in the hippocampus exhibit such a complex phenotype, where PSTUT is typically observed near the excitability level of a neuron. (e.g. CA1 neurogliaform [24], DG Total Molecular Layer subtype [15] etc.). Our simple models capture the characterizing features of both PSTUT and ASP. (Fig 2A) under the right input conditions. It is worth mentioning that all PSTUT neurons are inhibitory neurons and CA1 region has a proportionately larger number of these phenotypes [9, 25]. In many cases, however, the characteristic features of interrupted spiking can be only transiently present (Fig 2B). Here, a single pattern presents features of both bursting and spiking, where a relatively longer interval separates a few high frequency spikes (burst) from a train of regular spikes. In another set of examples, Granule cells and Hilar Ectopic Granule in the dentate gyrus (DG) show only transient bursting near excitability (Fig 2C and 2D). However, for an increased input current, Granule cells still showed the same class of TSWB.SLN with quantitative differences such as increased number of spikes, whereas Hilar Ectopic Granule transitioned to ASP. These constrained representations of two different DG neurons fall under the same family of non-persistent bursting, but they reflect finer quantitative differences in the input-dependent responses between these two neuron types. Thus, our simple models do not only qualitatively capture the rich diversity of dynamical classes defined systematically, but they are also quantitatively constrained representations of experimentally recorded patterns from the hippocampal neuron types.

### 3.2. Multi-compartment models as compact extensions of point-neuron models

The point-neurons presented in the last section would tremendously reduce the computational cost of simulating large-scale networks of hippocampal circuits. However, since they lack spatial dimension, they do not differentiate synaptic inputs from different layers, unlike their biological counterparts. For example, the hippocampal pyramidal neurons receive entorhinal projections on the apical dendrites in stratum lacunosum moleculare (SLM), and intra-hippocampal connections in stratum radiatum (SR), thereby compartmentalizing synaptic integration of distinct laminar inputs. While it is not possible to spatially segregate synaptic integration in a network of point-neurons, it is of interest to see the effects of such segregated synaptic integration mechanisms in a network. Hippocampome.org (ver 1.4) identifies 87 neuron types with their dendrites invading at least two layers. Therefore, for these neuron types, in addition to point neuron models, we created compact multi-compartment models with up to four compartments. Here, each compartment corresponds to a hippocampal layer, and this allows layer-level connectivity specifications for a neuron type.

One example for each of the four out of five possible multi-compartment layouts are illustrated here, and the fifth layout is discussed in detail in Section 3.3. The somatic compartment of a compact multi-compartment model quantitatively reproduces the spike patterns experimentally recorded from the soma of the respective neuron type for similar input currents (Fig 3 and Fig 4A). The number and layout of the coupled compartments is determined by the layers of dendritic invasion and known/possible soma locations of real neurons as illustrated by various examples in Fig 3. The dendritic compartments in a compact multi-compartment model are less excitable and have higher input resistances than the somatic compartment (Supplementary Fig S1B) [26, 27].

**Fig 3.**
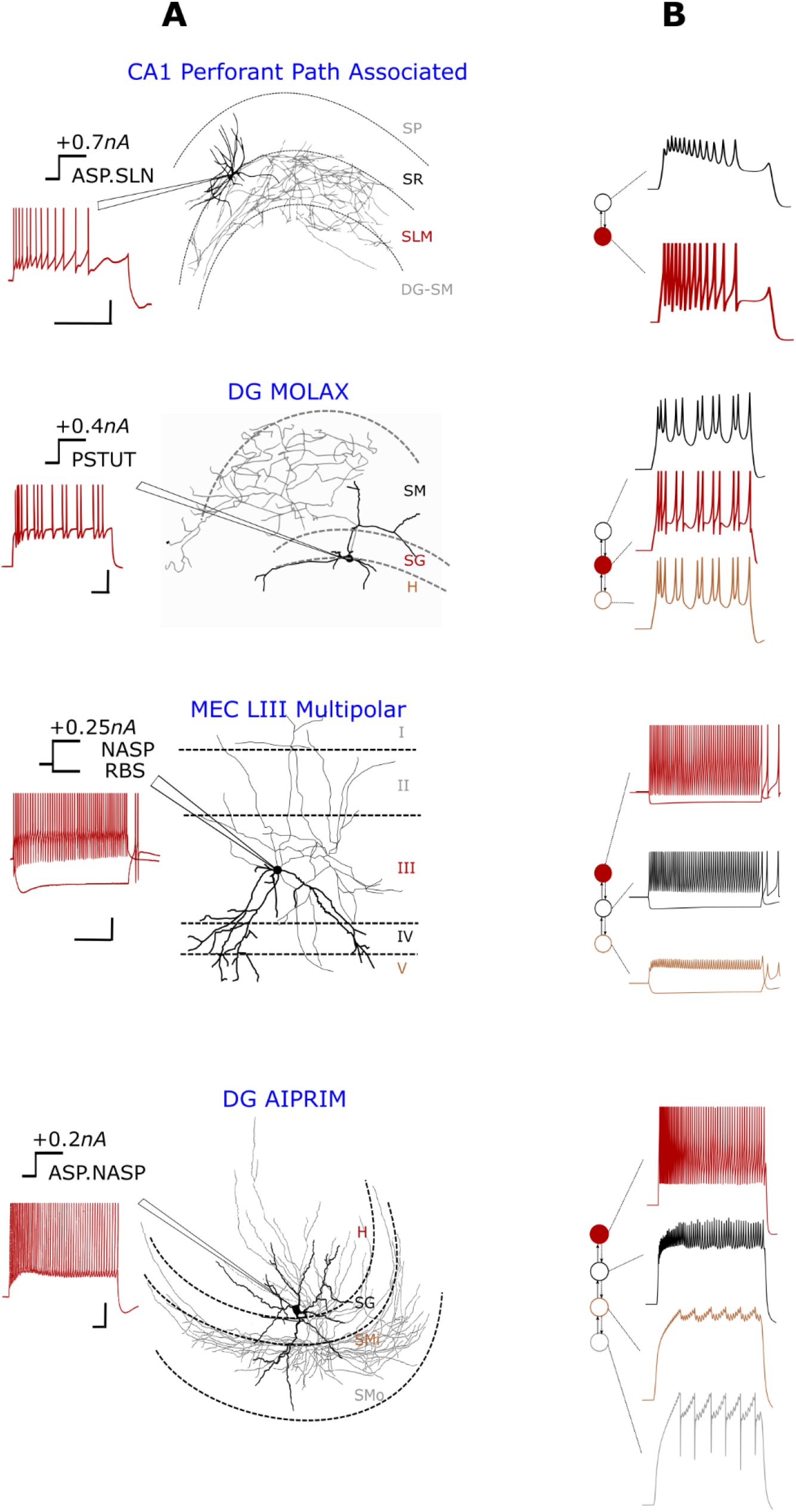
Multi-compartment models compactly extend point neurons to allow layer-level spatial context. (**A**) Experimentally recorded voltage traces (left) are given for four different morphological types (right). Dendritic invasion (darker) of layers and relative soma location determine the number and layout of compartments. (**B**) Layout of compartments coupled asymmetrically (left) correspond to the layers of dendritic invasion shown in A. Filled circles denote soma. Compartment responses for somatic input currents that are ±0.01*nA* from experimental input are given in right. See Fig 4 for quantitative comparison of spike pattern features, Supplementary Fig S1 for dendritic features, and Fig 5 for another possible 4-compartment layout. Experimental traces were digitized by *Hippocampome.org* from the following sources (from top to bottom): [28], [15], [18] and [29]. Morphological abbreviations: SMi and SMo – inner one-third and outer two-third of stratum moleculare. Experimental spike amplitudes are truncated. Calibration bars denote 200*ms*, 20*mV*.

**Fig 4.**
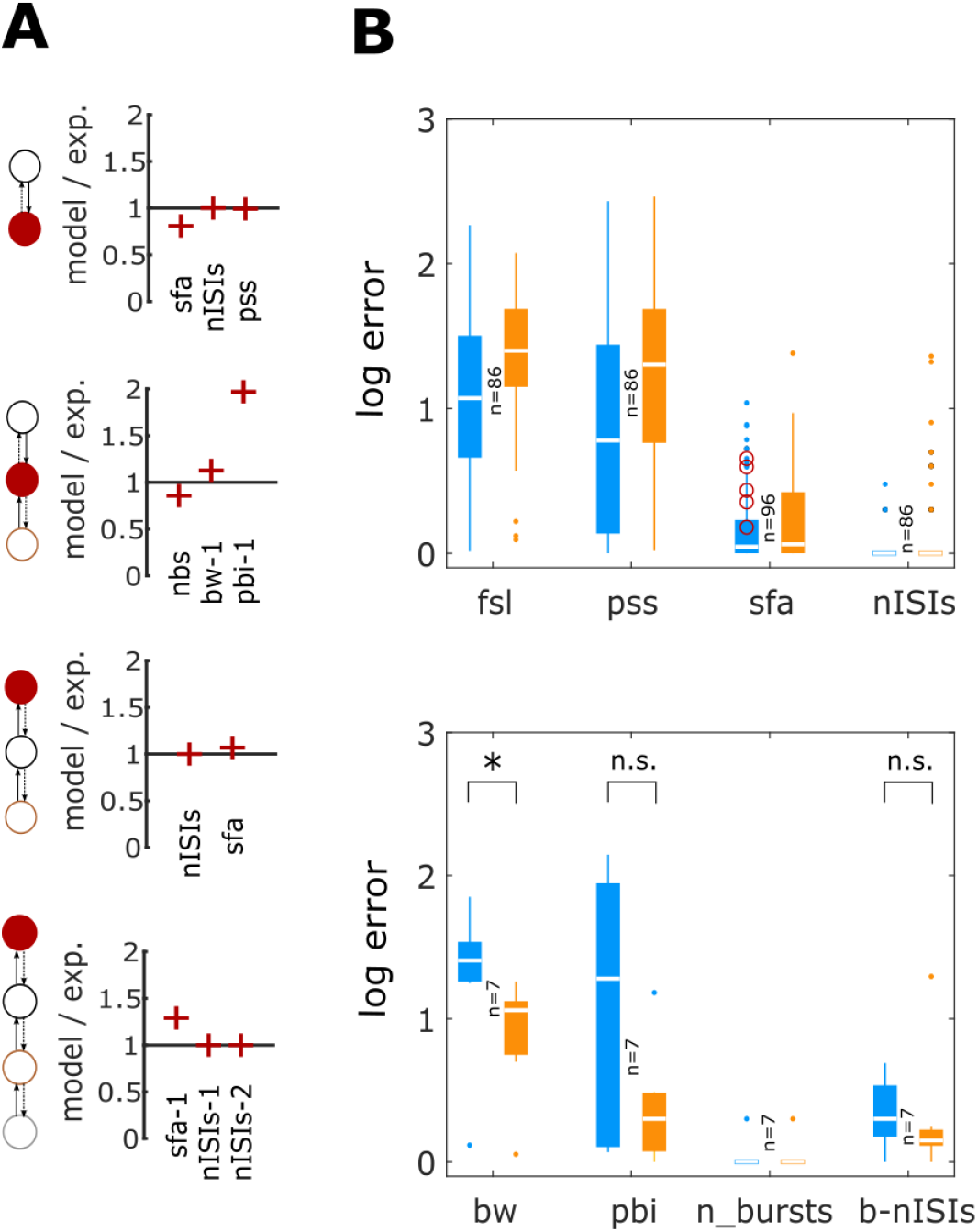
Accuracy of compact multi-compartment models in reproducing spike pattern features. (**A**) The quality of fit is given for key features as the ratio between simulated and experimental values for each of the four examples from Fig 3. (**B**) Pairwise comparisons of accuracy between single- (blue) and compact multi-compartment (orange) models for spiking features (top) and bursting features (bottom). While single-compartment models, in general, showed smaller errors for spiking features, they did not satisfy statistical criteria for RASP.ASP. patterns (denoted by circles in top panel). See Supplementary Fig S2 for an example for RASP.ASP. pattern. On the other hand, compact multi-compartment models generally improved the accuracy of bursting features (bottom) with a significant improvement in *bw* (p<0.005 for paired-sample *t*-test).

Furthermore, forward-coupling (from dendrite to soma) between compartments is just strong enough to evoke a somatic excitatory postsynaptic potential (EPSP) with an amplitude in the range [0.1, 0.9] *mV* for a single synaptic stimulation at a dendritic compartment and to achieve a forward-spike propagation (from dendrite to soma) ratio in the range [0.5, 1.0]. (Supplementary Fig S1C-D). As mentioned in methods, the backward-coupling (from soma to dendrite) is much stronger than the forward-coupling in most of our compact multi-compartment models, consistent with the electrotonic profiles reported for various neuron types [30–32]. Such an asymmetric design for coupling enables the somatic compartment to dominantly define the model’s overall intrinsic dynamics, while still preserving forward propagation properties for sub- and supra-threshold signals from dendrites. Thus, our multi-compartment models are compact extensions of point neuron models, which allow spatial contexts for synaptic integration (Fig S2).

Although the major motivation for creating compact multi-compartment models is to allow synaptic segregation in a network model, we also investigated if additional dendritic mechanisms implemented in our compact multi-compartment models could help achieve a better fitting of somatic spike patterns than their point-neuron counterparts. Therefore, we performed pairwise comparisons between the somatic spike pattern features of single-compartment and compact multi-compartment models. In general, implementing additional dendritic mechanisms in the models only improved the accuracy of bursting features (Fig 4B). Interestingly, *fsl* and *pss* errors were higher in the models due to the addition of dendritic compartments. However, it should be noted that each additional compartment not only adds two state variables, which require more computations for numerical simulation, but also adds ten open parameters (including coupling parameters) making it a more-challenging optimization task. It has been shown that adequate dendritic influence is necessary for bursting to exist in a 2-compartment model [12]. Although our single-compartment models were able to reproduce quantitatively comparable experimental bursting/stuttering patterns (Fig 2) (see [10] for two exceptions), compact multi-compartment models significantly improved the accuracy of *bw*, a key feature of bursting/stuttering patterns (Fig 4B).

Furthermore, while the single-compartment models quantitatively captured various classes of adapting spike pattern phenotype such as ASP., ASP.SLN, ASP.NASP and RASP.NASP, they failed to reproduce RASP.ASP. patterns. These patterns exhibit a strong and rapid adaptation in the first few ISIs, which is then followed by a very weak and sustained adaptation. Interestingly, we found that such a combination was not possible in the QM (red circles in Fig 4B), unless additional dendritic compartments were included. Two different time constants (parameter ‘*a*’) for the adaptation variable (state variable *U*) were required for the somatic- and dendritic-compartments respectively in order to capture such complex transients in the soma. In our single-compartment models, RASP.ASP. is represented by RASP.NASP, since the adaptation followed by RASP. is usually very weak. See Supplementary Fig S2 for an example.

### 3.3. Electrotonic compartmentalization in a 4-compartment model of CA1 pyramidal neurons

In addition to the features discussed in the last section, our compact multi-compartment models show electrotonic structures and interplay between different compartments that are similar to those experimentally observed in the pyramidal neurons of CA1. To illustrate this, here we present a 4-compartment model of CA1 pyramidal neurons and discuss the voltage attenuation and spike propagation properties of apical compartments. First of all, the somatic compartment captures the frequency adaptation (Fig 5B), the characterizing feature of the experimentally recorded spike pattern from a CA1 pyramidal neuron (Fig 5A), quantitatively (Fig 5C – left). Secondly, the dendritic compartments (SR, SLM and SO) are less excitable and have higher input resistances than the somatic compartment (Fig 5C – right).

**Fig 5.**
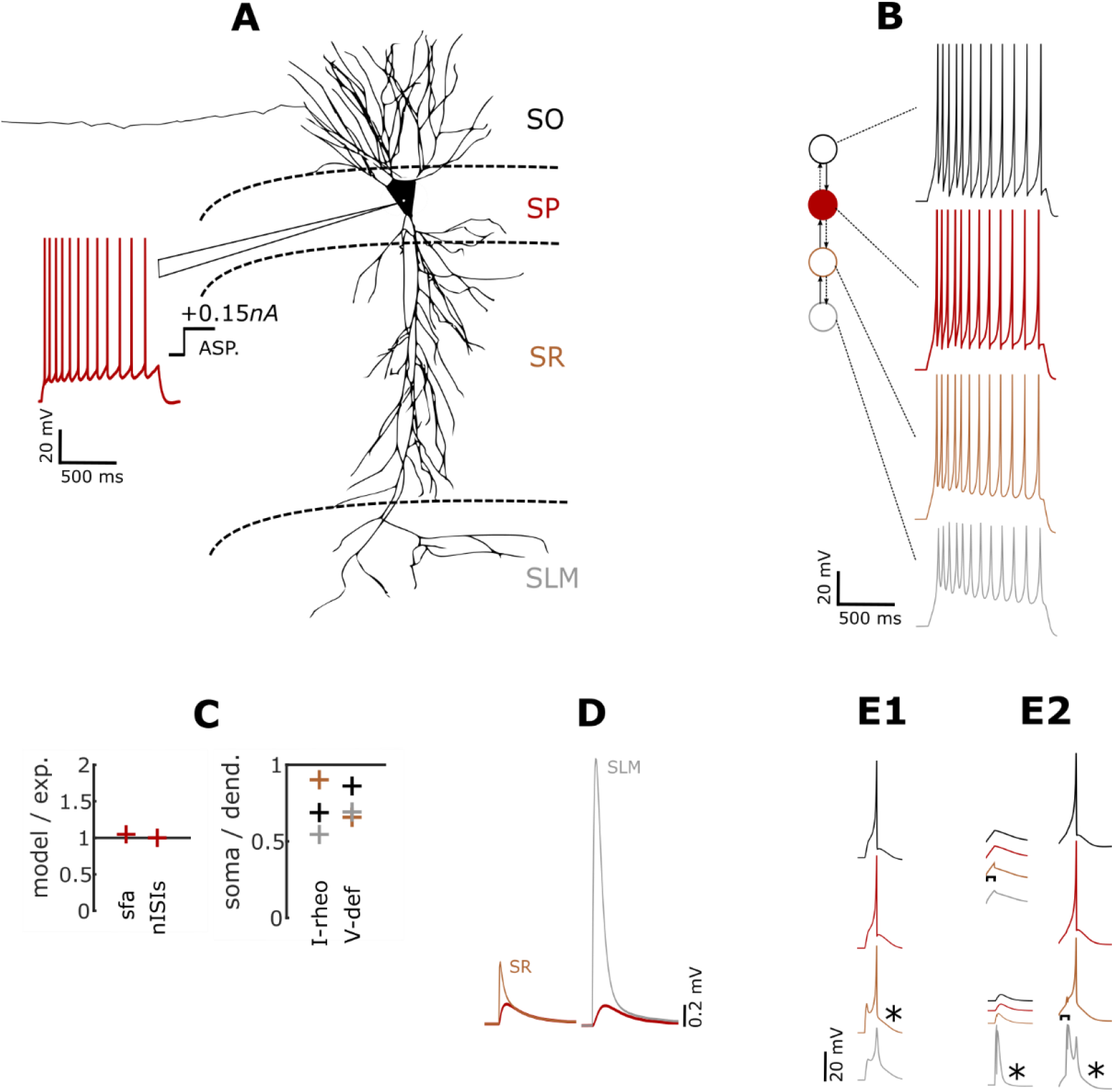
A 4-compartment model of CA1 Pyramidal neurons. (**A**) Experimentally recorded voltage trace from a CA1 Pyramidal neuron [33] digitized by *Hippocampome.org*, and digitally reconstructed morphology of the same type [34] reproduced from *Neuromorpho.org* [16]. (**B**) Layout of the compartments (left) and their responses to somatic input current with a magnitude of +0.14nA. (**C**) Somatic compartment reproduces key features that are quantitatively comparable to the experimental features (left). Minimum depolarizing input required to elicit a spike (I-rheo) and steady-state voltage deflection (V-def) for a hyperpolarizing input are higher in dendritic compartments than the somatic compartment. (**D**) Voltage attenuation from SLM to soma is much higher than the attenuation from SR in the model. (E1) A spike initiated at the compartment SR (denoted by ‘*’) successfully propagates to soma. (E2) A spike initiated at SLM failed to initiate a spike at soma (left-bottom), and additional depolarization level at SR using a small step current facilitates propagation of spike initiated at SLM (right).

In real neurons, integration of an EPSP is influenced by the location of the synapse, because the voltage attenuates more from a distal dendritic location to the soma, than from a proximal location. This is due to the higher input resistances of more distal dendrites with smaller diameters. However, it has been shown in some CA1 pyramidal neurons that the synapses might be able to compensate for their distance by scaling their conductances in order to sufficiently influence somatic voltage [35, 36]. In our model, compared to a synapse stimulated at SR to evoke a somatic (SP) EPSP with an amplitude of 0.2*mV*, a 12-fold increase in synaptic weight was required at SLM in order to evoke an EPSP with the same amplitude at SP (Fig 5D).

Furthermore, the distal compartments in our 3- and 4-compartment models rarely initiated a spike that successfully propagated to the soma. While the spike initiated at SR successfully triggered a somatic spike (Fig 5E1), the spike initiated at SLM failed to do so (Fig 5E2 – bottom in left). The same scenario, however, triggered a somatic spike, when SR was slightly depolarized further by a step current of small magnitude and duration (Fig 5E2 – right. Also see top in left). This is consistent with the experimental observation that the activation of CA1 neurons by the perforant path, which projects to SLM, is limited, and, modest activation of Schaffer-collateral synapses at SR facilitates forward propagation of distal spikes [37]. It has been suggested that Schaffer-collateral evoked EPSPs “gate” perforant path spikes in CA1 pyramidal neurons, and pyramidal neurons, in general, have functionally different dendritic domains [26]. To what extent these differences influence the emergent network properties, however, remains to be answered, and our models allow one to explore such questions.

### 3.4. Online repository of models: An enhancement to hippocampome.org

A comprehensive list of models of 68 types and 52 subtypes of neurons is freely available at *Hippocampome.org*. Mapping the intrinsic dynamics of each neuron type in a low-dimensional model space enhances the existing knowledge accumulated in this comprehensive knowledge base of hippocampal neuron types.

All the single-compartment and compact multi-compartment model parameters are presented in a matrix on the main page to enable easy browsing (Fig 6A). Within a neuron page, models for all subtypes (if any) for the given morphological type are available for download. This page includes both the experimentally recorded voltage traces and simulated ones for all subtypes (Fig 6B). Simulated spike patterns are also annotated with their class labels. Each type/subtype presents three downloadable files for the user (Fig 6C). A *Fit-file* includes both the experimental and simulated values for spike pattern features such as *fsl* and *sfa* for each available pattern in a JSON format. In addition, an XPP [38] script for single compartment models, and a csv input file that includes both single-compartment and compact multi-compartment models to be simulated using CARLsim [39], a high performance GPU-based simulator, are provided for each type. Links to help pages are provided under section “Simulation of Firing patterns” on *http://hippocampome.org* for model and feature description and instructions to run the scripts.

**Fig 6.**
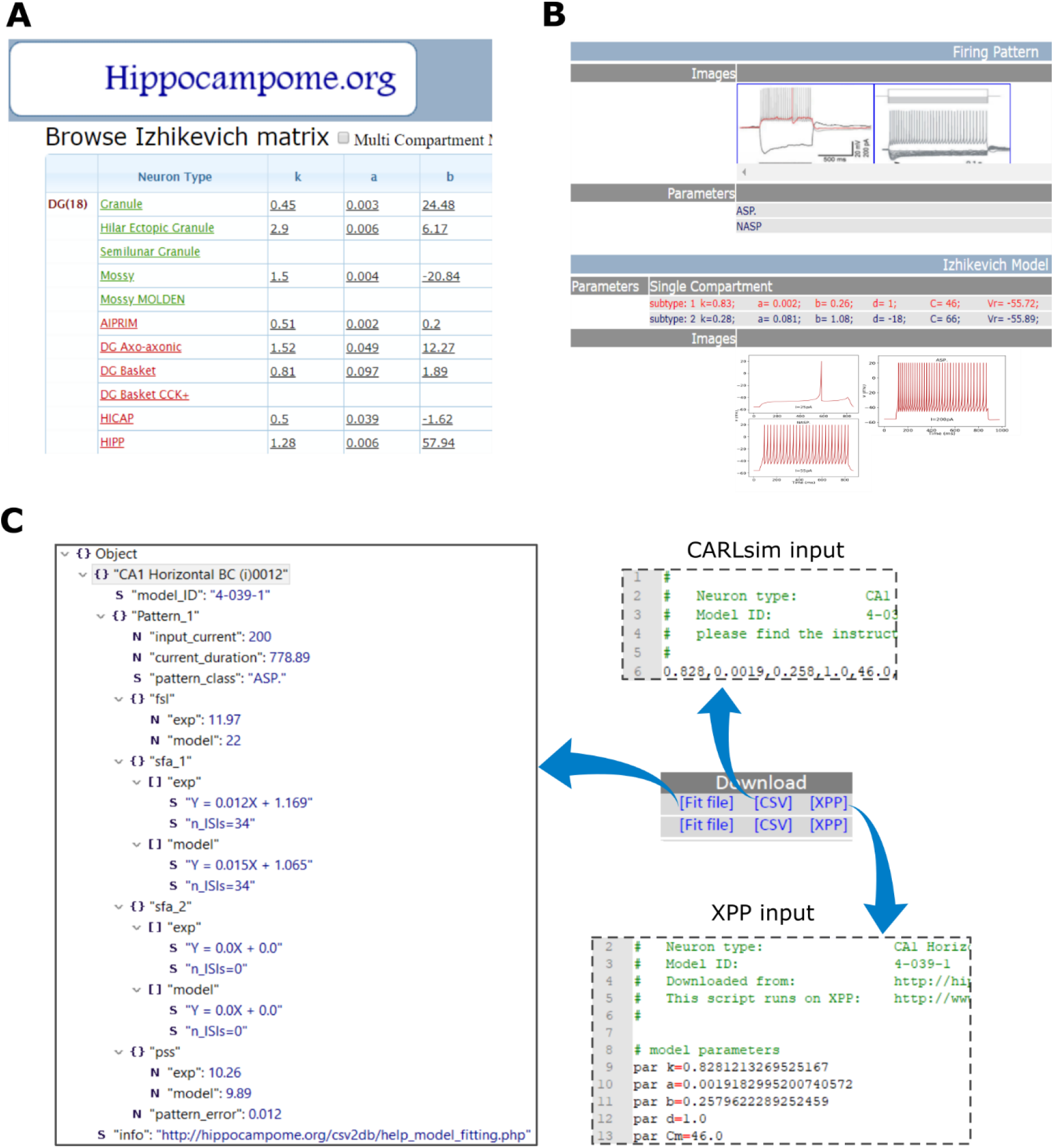
*Hippocampome.org* provides a comprehensive list of models and ready-to-run scripts. (**A**) Single- and multi-compartment model parameters for all neuron types are presented in a matrix form on the main page. Each row is linked to a neuron page (**B**) The neuron page for each neuron type has been enhanced to include model parameters and simulated traces for all types and subtypes (if any). (**C**) The neuron page provides the user with three downloadable files for each subtype: a *Fit-file* that lists both experimental and simulated features for each pattern, an XPP script to simulate single-compartment models, and a CARLsim input file for single- and multi-compartment models.

### 3.5. Relationship between model parameters and biological features

A limitation of the phenomenological model such as the QM used here is the lack of biological interpretability of its parameters. One advantage of current modeling work, which densely covers the diversity among neuron types, is that it allows one to explore relationships between the mathematical parameters of the QM and various known biological features. Our analysis revealed some interesting trends and correlations between the QM parameters and biological features, which are presented below.

In general, the parameters of the QM collectively determine its spike pattern phenotype. However, the parameter ‘*b*’, which determines if the model is an integrator (*b*<0) or a resonator (*b*>0), sufficiently distinguishes two families of phenotypes. Most of the models that show delayed spiking near their depolarizing excitability levels were found in the negative regions of ‘*b*’, whereas models that show rebound spiking for hyperpolarized input currents were strictly restricted to the positive regions (Fig 7A). Our results confirm the fact that all rebound spikers are resonators [11], and find that most delayed spikers are integrators with the exception of the ones found in the narrow range 0<*b*<20. Thus, rebound (Fig 1C) and delayed (Fig 1D) spiking are, in general, instances of two qualitatively very different types of intrinsic dynamics.

**Fig 7.**
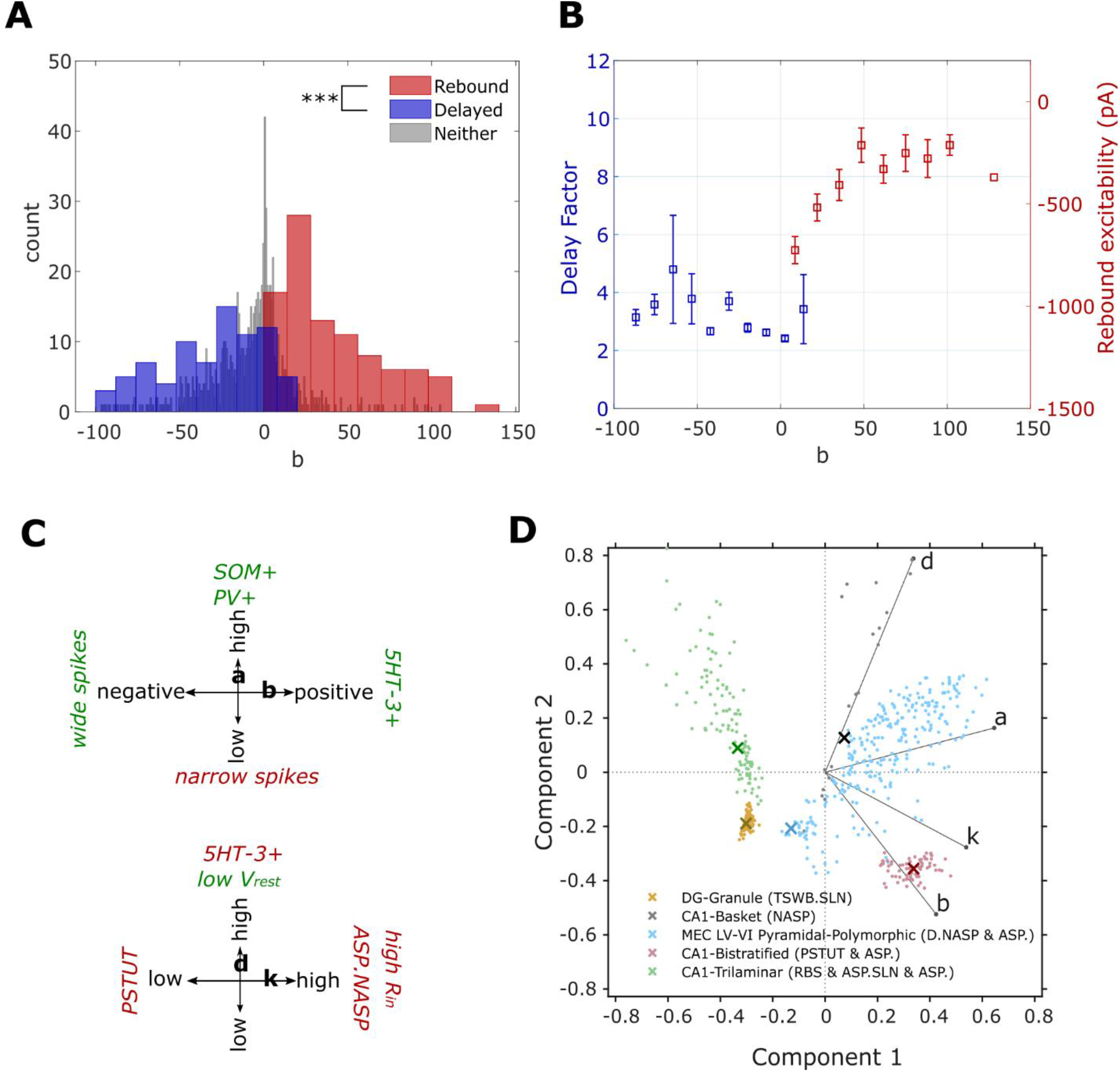
Relationship between model parameters and biological features. (**A**) Distribution of parameter ‘*b*’ for the models that show rebound spiking (red), delayed spiking (blue) and neither (grey). Rebound spiking types were strictly restricted to the positive region of ‘*b*’. In contrast, delayed spiking types were mostly found in the negative region (p<0.001 for two-sample t-test). (**B**) Mean and SEM of delay factors and rebound excitabilities of the models from the blue and red histogram bins respectively from A. Delay factor is the ratio between *fsl* and average of first two *ISIs*. Rebound excitability is the minimum magnitude hyperpolarizing current required to elicit rebound spikes. (**C**) Graphical representation of categorical correlations between model parameters and biological properties of neuron types (see BOX-1 for details). Green and red indicate the presence and absence respectively of a feature. (**D**) Separation of spike pattern phenotypes in the space of first two principal components. ‘x’ denotes the best model for each type. Vectors denote the principal component coefficients of the respective parameters.

Next, we studied how much the parameter ‘*b*’ quantitatively influences the respective features in delayed and rebound spiking types. We used delay factors for the former and measured rebound excitability levels for the latter. Increasing ‘*b*’ makes the model more rebound excitable until ‘*b*’ reached a value of +50, beyond which there was no noticeable effect (Fig 7B). Furthermore, there was no clear trend in the relationship between ‘*b*’ and delay factor. Thus, while ‘*b*’ alone can define a sharp qualitative change in the intrinsic dynamics, its interaction with other parameters such as ‘*a*’ determine precise quantitative features.

In addition, several interesting trends between the model parameters and electrophysiological and molecular properties of neuron types were revealed in pairwise correlations. For instance, stuttering phenotype (PSTUT) was never found with low values (bottom-third) of ‘*k*’, and high values (top-third) of input resistance was never found with high values (top-third) of ‘*k*’. This is consistent with a correlation reported previously that all PSTUT neuron types have either low or intermediate input resistances [9]. See Box-1 for more of such relationships and Fig 7C for a graphical representation.

#### Box 1.

Categorical correlations between model parameters and electrophysiological and molecular properties in hippocampal neurons

1. None of the neuron types that show **PSTUT** has a low value for ***‘k’*** (p<0.005, n=55). In contrast, none of the neuron types that show ASP.NASP has a high value for ***‘k’*** (p<0.005, n=55). Moreover, no neuron types with high input resistance (***R_in_***) has a high value for ***‘k’*** (p<0.001, n=23).
2. None of the 26 neuron types except CA3 Lucidum ORAX has **narrow spikes** and a low value for ***‘a’*** (p<0.005). Moreover, clearly positive expressions of somatostatin (**SOM**) or parvalbumin (**PV**) tend to co-occur with high values of ***‘a’*** (p<0.01, n=24 and p<0.05, n=44 respectively).
3. Neuron types with **wide spikes** tend to have negative values for ***‘b’*** (p<0.001, n=37). In contrast, positive expression of serotonin (5HT-3) co-occurs with positive values for ***‘b’*** (p<0.05, n=31).
4. Low values of resting voltage (***V_rest_***) tend to co-occur with high values of ***‘d’*** (p<0.01, n=27). In contrast, no neuron type with positive expression of serotonin (**5HT-3**) has a high value for ***‘d’*** (p<0.05, n=23).

The p values and sample sizes (n) pertain to Barnard’s exact test for 2 × 2 contingency tables (see ‘Materials and methods’).

Finally, our modeling framework represents each neuron type as a cloud of possibilities in the model parameter space (Fig 7D). Spike patterns produced by all the models in a cloud strictly adhere to the criteria for the respective target qualitative class, but small errors in the quantitative features were accepted to allow variabilities in the spike pattern features (not shown here). See [10] for more details on the optimization framework design that allows such variabilities and some examples of ranges of quantitative features. At present, these clouds are only identified for single-compartment models due to the computational cost of exploring higher dimensional parameter spaces of multi-compartment models.

## 4. Discussion

The simple models presented in this work are aimed at creating large-scale network models of hippocampal circuits that are biologically realistic, yet computationally efficient. We first discuss biological realism in the context of variability in the intrinsic dynamics and then discuss how one can take advantage of the computational efficiency of these models in creating network models.

Hippocampal neurons show diverse features in their morphological, electrical and molecular properties. Hippocampome.org (ver 1.4) identifies 122 types of neurons defined primarily based on their neurite invasion patterns in the hippocampal parcels [8]. Their intrinsic spike pattern features were extracted from relevant publications, and systematic characterization of such features revealed diverse and complex spike pattern phenotypes among the 122 morphological types [9]. Current work presents a comprehensive set of simple models that are accurate quantitative representations of such spike pattern phenotypes. However, it is worth discussing their accuracy in a broader context. The intrinsic property of a neuron revealed in its spike patterns is determined by the types and precise distribution of the underlying ion channel conductances such as sodium, potassium, and calcium. However, it has been shown that similar dynamics can arise from a broad range of combinations of these conductances [40–42]. Consistent with this notion, our modeling framework represents a spike pattern phenotype as a cloud of possibilities in the parameter space (Fig 7D). Two closely related issues motivate such representation.

First issue is the existence of intrinsic variabilities in the spike pattern features among different neurons of the same type. For example, all the models representing CA1 Trilaminar type (Fig 7D) were obtained using the features of voltage traces recorded from a single neuron (Fig 1C). While this particular neuron elicited 22 spikes with an *sfa* magnitude of 0.038 for 0.05*nA*, a different CA1 Trilaminar neuron might show slightly different values for these features under the same input conditions due to intrinsic variability. Furthermore, the recorded intrinsic spike pattern features might be influenced by the conditions such as the type of recording electrode and difference in the species. However, current knowledge about the intrinsic dynamics of these neuron types is limited to the representative traces that the researchers who studied these neuron types chose to publish. Therefore, we allowed a small range in the spike pattern features of a model as long as these features strictly adhere to the definitions of the respective target qualitative class. While the cloud boundaries defining such ranges are currently arbitrary, one could easily enhance our modeling framework to include more realistic ranges, when such ranges are experimentally obtained for all neuron types.

Secondly, neurons have intrinsic plasticity and undergo homeostatic regulations to maintain some constancy in the network activity [42–46]. In cell cultures, intrinsic homeostasis has been shown to modify pharmacologically isolated neurons’ non-synaptic ion channel conductances. Such modifications shift the input-dependency of a neuron’s responses based on the history of activity. For example, activity deprived neurons showed higher firing rates than control group for the same magnitude current injections [46]. In another study, chronic isolation from normal inputs switched a neuron’s response from tonic spiking to intrinsic bursting and this transition was reversed by applying a rhythmic inhibitory drive [43]. While these results suggest that each neuron has a working range that flexibly defines its input-dependent responses, such ranges likely preserve the overall qualitative spike pattern phenotypes [45]. Our EA search for a cloud of models not only included the space of intrinsic QM parameters that define a phenotype, but also included a small range for input current (a 20pA range symmetrically encompassing experimental input current magnitude), allowing a little flexibility for its input-dependency.

Considering the issues discussed above, an approach to modeling biological circuits should assume a flexible range for its components. While Hebbian plasticity rules can enable flexible ranges in synaptic conductances, the rules governing a neuron’s intrinsic plasticity remain largely unknown. Although cell-autonomous regulatory rules have been proposed [47], from a network perspective, intrinsic homeostasis have been shown to synergistically result from multiple interacting components in a circuit [48, 49]. Exhaustively reductionist approaches to modeling brain regions specify *precise* descriptions at the level of ion channel conductances. While data gathered from different experimental conditions or inevitably from different animals drive such intrinsic descriptions, there is no guarantee that they specify dynamically compatible critical ranges necessary for a higher-level integrative property [50].

A large-scale approach to modeling a brain region, rather than being purely reductionist, should attempt to complement the descriptions of individual components with syntactically relevant descriptions at integrative level. For example, temporal sequences of activity in ensembles of hippocampal neurons are correlated with the locations of an animal during spatial navigation [51–53]. Such self-organizing ensembles of neurons, in general, have been suggested to form neural syntax [54]. Complex periodic structures in these ensembles such as theta-modulated gamma activity patterns should be enforced in a network model as sparse higher-level descriptions.

Future studies should aim to identify a family of models for an experimentally known network-level property within the anatomical constraints of connectivity among hippocampal neuron types [55] using the sampling regions for those types created in this study. Then, the identified family of models should be evaluated for their predictive power, or one could investigate how the predictive abilities increase by scaling up the network, or by adding more mechanisms such as synaptic plasticity and spatial context for synaptic integration. This approach emphasizes the goal of creating the simplest model with the most predictive power iteratively.

Finally, it is important to identify recurring patterns of self-organization in biological complex systems and translate such patterns into mathematical descriptions that could be enforced on a meta-level using self-adaptive techniques such as an evolutionary algorithm that heuristically explores the given parameter space. If a biological complex system can indeed allow a little flexibility and compensation among multi-level components, then it suggests that a certain property could emerge from multiple, similar configurations in a network parameter space, which a metaheuristic approach can take advantage of. While this might be a computationally expensive task, our simple models with only two state variables per neuron as opposed to hundreds in a biophysically detailed multi-compartment model allow one to approach this problem much more efficiently. Future releases of *Hippocampome.org* are aimed at approximating the counts of different neuron types and mapping synaptic properties to potential connections. These enhancements will further narrow down the space of biological possibilities to create realistic large-scale models of hippocampal circuits.

## Acknowledgements

This work is supported by grants R01NS39600 (NIH) and U01MH114829 (NIH). We thank Abhijeet Mishra and Nikhil Koneru for helping with the implementation of portal features, and Ernest Barreto and Keivan Moradi for useful discussion.

