## Supplementary material for "Simple models of quantitative firing phenotypes in hippocampal neurons: comprehensive coverage of intrinsic diversity"

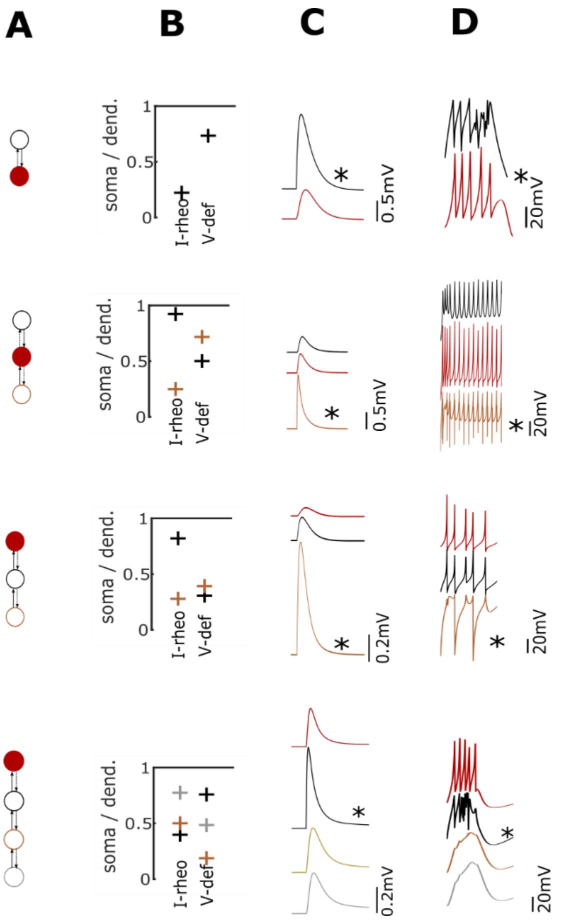


S1 Fig. **Multi-compartment models capture qualitative dendritic properties and sub- and supra-threshold signal propagation.** (A) Four different layouts of asymmetrically coupled compartments from Fig 3. (B) Minimum depolarizing input required to elicit a spike (I-rheo) and steady-state voltage deflection (V-def) for a hyperpolarizing input are higher in dendritic-compartments than the somatic-compartment. (C) A single synapse stimulated at a dendritic-compartment (denoted by '*') evokes an unitary EPSP at the somatic-compartment (red traces) with an amplitude in the range [0.1, 0.9] mV. (D) Coupling mechanism implemented in the models allows forward propagation of spikes initiated at a dendritic-compartment (denoted by '*').

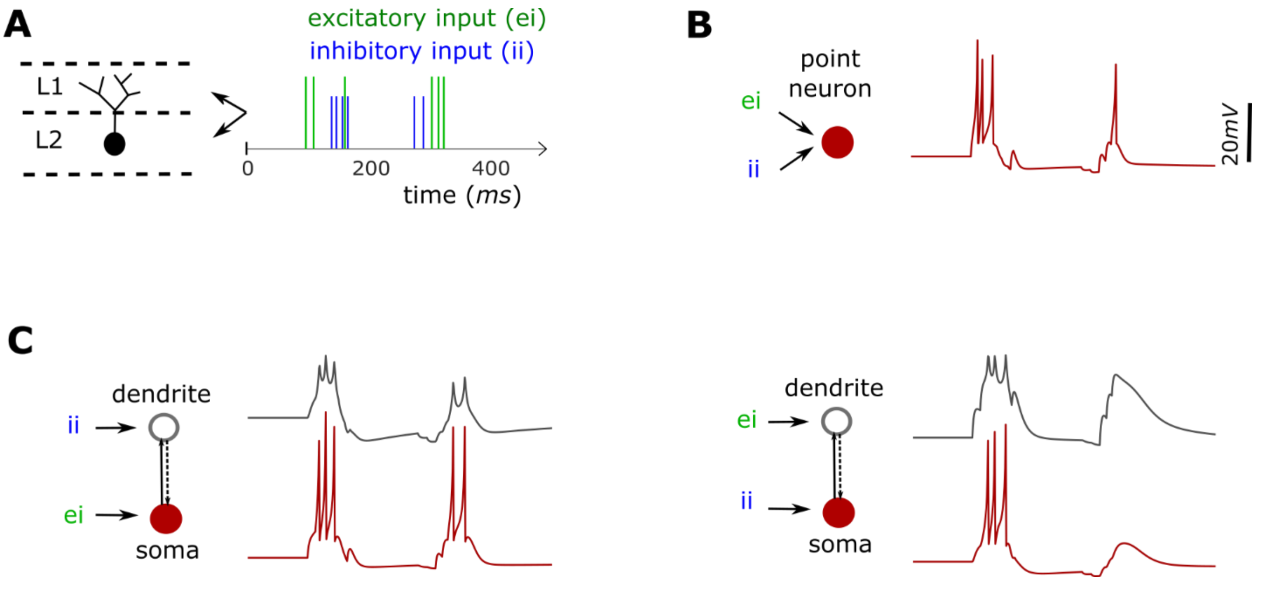


S2 Fig. **Multi-compartment models allow spatial context for integration of presynaptic spikes.** (**A**) A schematic illustration of a biological neuron with its soma and dendrites in different layers L1 and L2. This neuron receives two distinct presynaptic spike trains in L1 and L2. (**B**) A simplified point-neuron model integrates both the excitatory and inhibitory presynaptic spikes at the same point. (**C**) A 2-compartment model (see Fig 3) can integrate distinct inputs in different compartments. The model behaves differently (left vs. right) depending on the location of integration of distinct inputs.

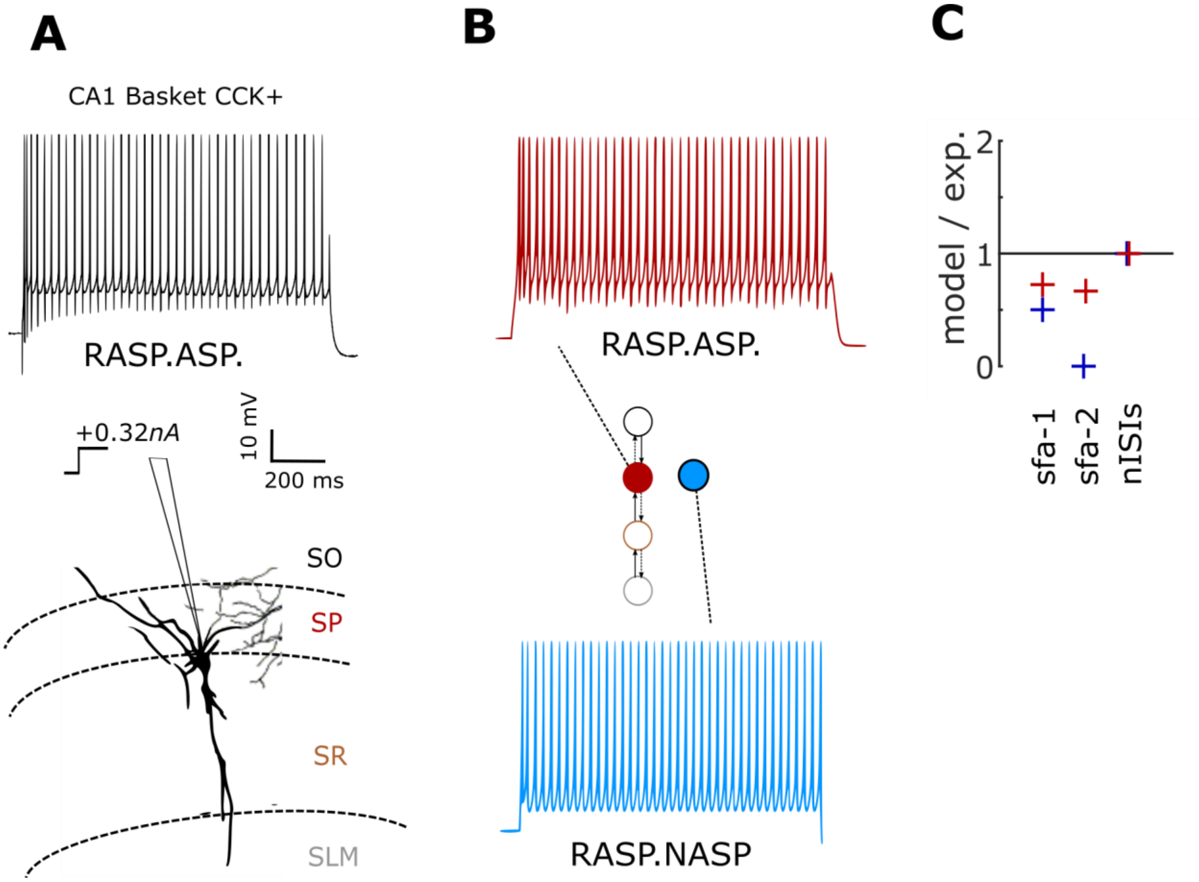


S3 Fig. **Additional compartments are necessary to capture the complex transient pattern RASP.ASP.** (A) Experimentally recorded voltage trace from a CA1 Basket CCK+ neuron (Cope et al. 2002) digitized by Hippocampome.org. (B) 4-compartment model reproduces the pattern RASP.ASP. (red), and the single-compartment counterpart failed to do so (blue). (C) While both versions reproduce nISIs accurately, the multi-compartment model more accurately reproduces sfa. Sfa-1 is the rapid frequency adaptation measured in the first three ISIs (RASP.), and sfa-2 is the weak adaptation measured in the remaining 35 ISIs (ASP.) Note that sfa-2=0 in the single-compartment model. Spike amplitudes are truncated.
